# The HLA-II immunopeptidome of SARS-CoV-2

**DOI:** 10.1101/2023.05.26.542482

**Authors:** Shira Weingarten-Gabbay, Da-Yuan Chen, Siranush Sarkizova, Hannah B. Taylor, Matteo Gentili, Leah R. Pearlman, Matthew R. Bauer, Charles M. Rice, Karl R. Clauser, Nir Hacohen, Steven A. Carr, Jennifer G. Abelin, Mohsan Saeed, Pardis C. Sabeti

**Author notes:** Corresponding author (S.W-G). These authors contributed equally (Shira Weingarten-Gabbay, Da-Yuan Chen, Siranush Sarkizova). Senior author (Jennifer G. Abelin, Mohsan Saeed, Pardis C. Sabeti).

## Abstract

Targeted synthetic vaccines have the potential to transform our response to viral outbreaks; yet the design of these vaccines requires a comprehensive knowledge of viral immunogens, including T-cell epitopes. Having previously mapped the SARS-CoV-2 HLA-I landscape, here we report viral peptides that are naturally processed and loaded onto HLA-II complexes in infected cells. We identified over 500 unique viral peptides from canonical proteins, as well as from overlapping internal open reading frames (ORFs), revealing, for the first time, the contribution of internal ORFs to the HLA-II peptide repertoire. Most HLA-II peptides co-localized with the known CD4+ T cell epitopes in COVID-19 patients. We also observed that two reported immunodominant regions in the SARS-CoV-2 membrane protein are formed at the level of HLA-II presentation. Overall, our analyses show that HLA-I and HLA-II pathways target distinct viral proteins, with the structural proteins accounting for most of the HLA-II peptidome and non-structural and non-canonical proteins accounting for the majority of the HLA-I peptidome. These findings highlight the need for a vaccine design that incorporates multiple viral elements harboring CD4+ and CD8+ T cell epitopes to maximize the vaccine effectiveness.

## INTRODUCTION

Recent success with synthetic vaccines against viral diseases has demonstrated their promise for future use, but also highlighted the need for a deeper understanding of how the adaptive immune system recognizes and responds to viral infections(Sette & Crotty, 2021; Wherry & Barouch, 2022). In contrast to traditional attenuated vaccines that mimic the natural interaction between viruses and the immune system, rationally designed synthetic vaccines deliver only a small, selected portion of the viral genome, often a single protein, to stimulate an immune response. This allows rationally designed vaccines to be developed quickly, produced in large quantities with low-cost manufacturing, and administered safely (Pardi et al., 2018). However, the development of rationally designed vaccines necessitates a comprehensive knowledge of viral immunology and the parts of the viral genome that are recognized by the different arms of the immune system.

Limitations of the COVID-19 vaccines developed during the course of the pandemic highlighted the importance of eliciting a more comprehensive immune response through T cells(Neale et al., 2023). Early vaccines mostly focused on eliciting humoral immune response through utilizing the viral spike protein (S) (Baden et al., 2021; Martínez-Flores et al., 2021; Sahin et al., 2021). However, challenges with waning immunity led to a greater focus on providing durable immunity by eliciting CD8+ cytotoxic and CD4+ helper T cell responses. Together with the high frequency of mutations in spike and the associated immune evasion of emerging variants, these findings motivated researchers to incorporate T cell targets, such as the nucleocapsid (N), in the next generation of COVID-19 vaccines (Arieta et al., 2023; Dutta Noton K. et al., 2020; Hajnik et al., 2022; Matchett et al., 2021; Oronsky et al., 2022). CD8+ and CD4+ T cells recognize viral peptides that are endogenously processed and presented on the surface via HLA-I and HLA-II complexes, respectively. These T cell epitopes originate from all viral proteins, in contrast to neutralizing antibodies that are mostly confined to external structural viral proteins. Although the presence of a large number of potential T cell epitopes in the viral genome offer a wide range of candidates, it can also present a challenge in identifying the most effective targets for T cell-based vaccines.

To understand the full range of T cell epitopes, it is important to implement non-targeted, comprehensive approaches in addition to conventional T cell assays. Targeted T cell assays require researchers to decide a priori which peptides to screen in the experiment, based on assumptions about which viral proteins are expressed and processed for HLA presentation. While T cell responses to SARS-CoV-2 have been extensively studied, many studies have focused on only a subset of viral proteins, particularly the structural proteins S, N, membrane (M), and envelope (E), rather than considering the full range of proteins in the SARS-CoV-2 proteome (Chen et al., 2021; Joag et al., 2021; Keller et al., 2020; Le Bert et al., 2021; Lee et al., 2021; Mahajan et al., 2021; Nielsen et al., 2021; Rha et al., 2021; Shomuradova et al., 2020). A smaller number of studies have looked at T cell responses to peptides from all annotated viral proteins and found significant responses to both structural and non-structural proteins (Ferretti et al., 2020; Gangaev et al., 2021; Kared et al., 2021; Mateus et al., 2020; Nelde et al., 2021; Prakash et al., 2021; Tarke et al., 2021). Although these studies provide a broader view, they have still generally only included peptides from annotated canonical proteins and have not considered peptides from non-canonical ORFs that were identified through experimental translation measurements (Finkel et al., 2020).

Taking an untargeted approach of HLA immunopeptidome profiling, we recently revealed surprising insights about HLA-I presentation in SARS-CoV-2 infected cells. Immunopeptidome profiling utilizes mass spectrometry to detect peptides that are endogenously presented on the HLA complex in different disease contexts (Abelin et al., 2017; Bassani-Sternberg & Gfeller, 2016; Chong et al., 2018; Croft et al., 2013; McMurtrey et al., 2008; Rucevic et al., 2016; Sarkizova et al., 2020; Schellens et al., 2015; Ternette et al., 2016). By applying it to SARS-CoV-2 infected cells we uncover HLA-I peptides from canonical proteins as well as overlapping ORFs in the coding region of N and S that were overlooked by previous T cell studies (Weingarten-Gabbay et al., 2021). Strikingly, some of the non-canonical peptides were more immunogenic in COVID-19 patients and humanized mice when compared to the strongest canonical protein derived antigens reported to date. Two of the non-canonical peptides were further supported by an independent HLA-I peptidome study from another group (Nagler et al., 2021). We also uncovered that, in contrast to earlier studies focusing mostly on structural proteins, non-structural proteins constitute a significant portion of the HLA-I peptidome, and that the timing of viral protein expression is a key determinant of HLA-I presentation and immunogenicity. The list of SARS-CoV-2 HLA-I peptides resulting from our study was recently used to characterize a T cell-directed mRNA vaccine (BNT162b4) (Arieta et al., 2023). Of the 19 HLA-I peptides observed in cells expressing a multi epitope vaccine, 3 were identical to peptides that we detected in infected cells. Together, HLA immunopeptidome profiling can identify new, highly potent T cell epitopes, inform vaccine design, and deepen our understanding of the determinants of viral antigen presentation.

Immunopeptidome profiling can be utilized to directly characterize another critical process in the antiviral immune response: HLA-II presentation to CD4+ helper T cells. In contrast to HLA-I presentation that samples endogenous cytosolic proteins, HLA-II presents peptides from proteins that have been taken up from outside the cell via endocytosis. Hence, different viral proteins may differentially access the HLA-I and HLA-II pathways. Understanding these differences can enable researchers to design vaccines that elicit both CD8+ and CD4+ T cells responses. Previous HLA-II peptidome studies of SARS-CoV-2 researched HLA-II peptides derived only from the spike protein, using a purified recombinant protein (Knierman et al., 2020; Parker et al., 2021), or from four viral proteins (N, M, E and nsp6) using plasmid overexpression (Nagler et al., 2021). Thus, we have not yet achieved a systematic view of HLA-II peptides from the full SARS-CoV-2 genome in the context of authentic virus infection.

In this study, we set out to achieve a comprehensive map of SARS-CoV-2 peptides that are processed and presented by HLA-II complexes. Using these data, we dissected the source viral proteins and the processed regions within each viral protein that are presented by HLA-II. We contextualize these findings by using thousands of reported CD4+ T cell epitopes, inferring the contribution of antigen processing and presentation steps to T cell responses observed in COVID-19 patients. We then compared the immunopeptidome of HLA-I and HLA-II complexes, revealing important differences between SARS-CoV-2 presentation to CD4+ and CD8+ T cells. The newly identified CD4+ T cell targets and the insights our study rendered about viral HLA-II presentation will enable a more precise selection of peptides for the next generation of COVID-19 vaccines that aim to target multiple viral proteins. The concepts learned from this study can also be applied to the design of synthetic vaccines against other viral pathogens.

## RESULTS

### Immunopeptidome profiling of HLA-II peptides in SARS-CoV-2 infected cells

To interrogate the HLA-II immunopeptidome of SARS-CoV-2, we infected A549 and HEK293T cells with the virus after inducing the HLA-II presentation pathway, immunoprecipitated (IP) HLA-lI-peptide complexes, and identified bound peptides by liquid chromatography-tandem mass spectrometry (LC-MS/MS)(**Figure 1A**). Both cell lines stably expressed ACE2 and TMPRSS2, two important SARS-CoV-2 entry factors. To induce the HLA-II pathway, we transduced the cells with the MHC class II transactivator (CIITA) using a lentiviral vector. Overexpression of CIITA, a master transcriptional regulator, facilitates cell-surface expression of HLA-II complexes and the peptide-loading machinery and has been previously used to interrogate the HLA-II immunopeptidome of virus-infected cells (Becerra-Artiles et al., 2019, 2022) and tumors (Forlani et al., 2021; Hos et al., 2022). In addition, some lung epithelial cells express HLA-II (Neuwelt et al., 2020; Wosen et al., 2018) and thus, studying the HLA-II immunopeptidome of infected cells mimics HLA-II presentation that can occur *in-vivo* during the course of infection with a respiratory virus.

**Figure 1.**
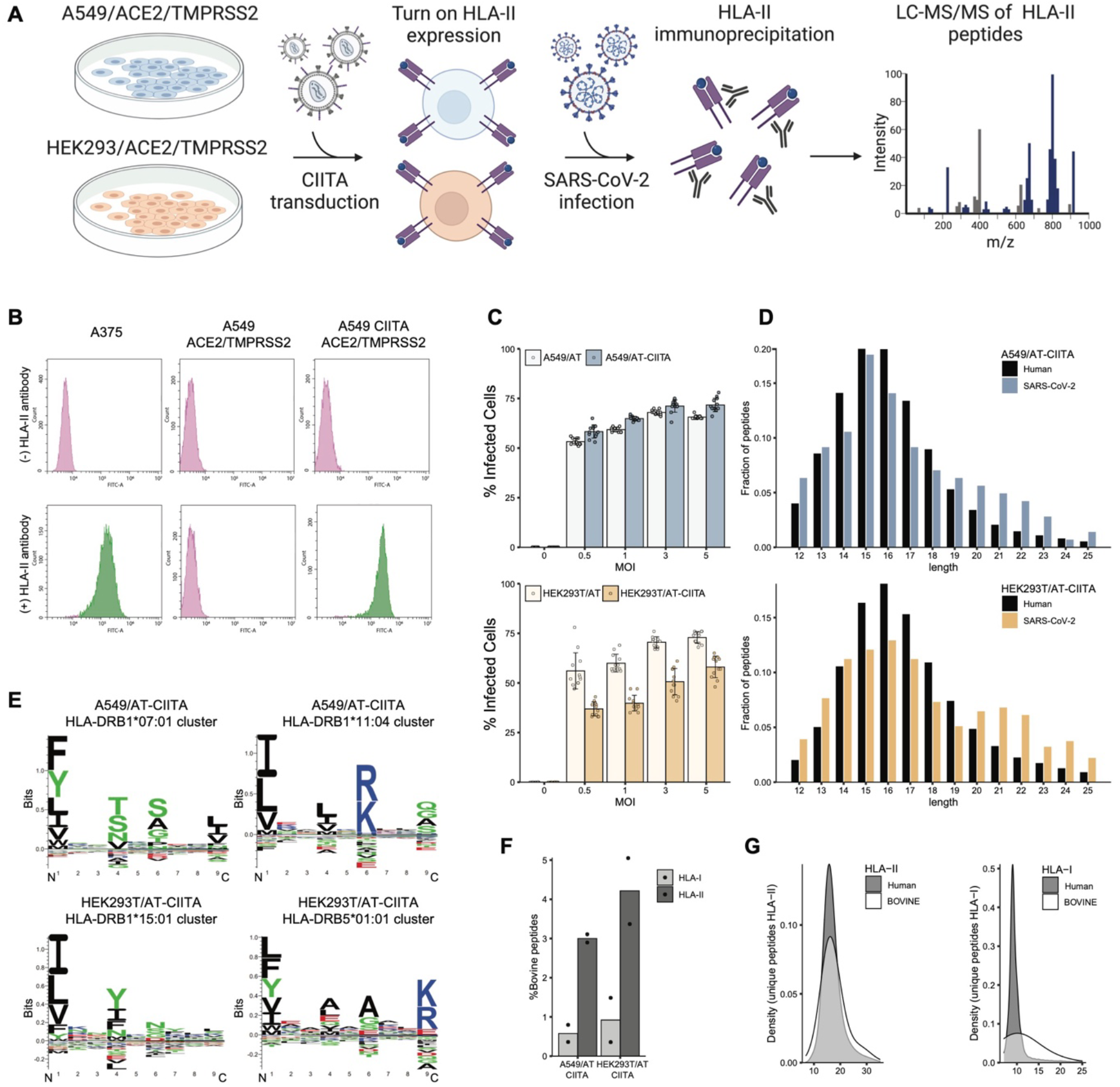
HLA-II immunopeptidome profiling of SARS-CoV-2 infected cells. **(A)** Schematic of the experimental workflow. **(B)** HLA-II expression measured by flow cytometry using a FITC-conjugated anti HLA-DR/DP/DQ antibody. Gating was based on unstained cells. **(C)** SARS-CoV-2 infection levels in A549/AT and HEK293T/AT cells with and without CIITA transduction. The infection was quantified using immunofluorescence staining for the nucleocapsid protein. MOI = Multiplicity Of Infection. **(D)** Length distribution of the eluted peptides in SARS-CoV-2 infected cells. **(E)** Gibbs Cluster deconvoluted motifs of the eluted peptides. **(F)** The percentage of peptides originating from Bovine proteins detected in the HLA-I and HLA-II immunopeptidomes. HLA-I data are from Weingarten-Gabbay et al. (Weingarten-Gabbay et al., 2021) **(G)** Length distribution of human and bovine peptides in the A549 and HEK293T HLA-I and the HLA-II immunopeptidomes.

We characterized the CIITA-transduced cells to ensure proper induction of proteins in the HLA-II pathway. First, we compared cell surface levels of HLA-II in A549 cells with those in a positive control human melanoma A375 cell line that endogenously expresses HLA-II (Deffrennes et al., 2001). The cell surface flow cytometry revealed strong induction (∼80-fold) of HLA-II expression in A549/ACE2/TMPRSS2 (A549/AT) cells upon CIITA transduction, with a similar fluorescence intensity as in A375 cells (**Figure 1B**). To monitor the expression of additional proteins in the HLA-II pathway, we examined the whole proteome of A549/AT and HEK293T/AT cells by analyzing lysates after the HLA-II IP using LC-MS/MS. We observed the expected increase in CIITA-induced proteins that are localized in the MHC-II region of the MHC locus including HLA-DM, HLA-DO, and TAP1 (**Figure S1A**).

We also ensured that CIITA overexpression did not affect the cell susceptibility to SARS-CoV-2 infection, since a recent study reported that CIITA can restrict SARS-like coronaviruses in U2OS osteosarcoma cells (Bruchez et al. 2020). We quantified SARS-CoV-2 infection by immunofluorescence (IF) staining of cells with an anti-N antibody at 24 hours post infection (hpi) (**Figure 1C**, **Figure S1B**). When infected at different multiplicities of infection (MOIs), and A549/AT cells exhibited similar infection levels regardless of CIITA expression. In contrast, HEK293T/AT cells showed reduced infection upon CIITA expression, although a substantial number of cells were still positive, with ∼50% infected cells at MOI of 3 (the MOI we used for our HLA-II IP experiment).

To gauge the technical performance of our experimental system, we examined if the peptides detected by LC-MS/MS match known characteristics of HLA-II peptides. We performed HLA-II IP of non-infected and infected cells in two biological replicates at 24 hpi using a mixture of antibodies targeting the three HLA-II loci (HLA-DR, HLA-DP, and HLA-DQ). We recovered 21,541 and 29,600 unique HLA-II peptides from A549/ACE2/TMPRSS2/CIITA (A549/ATC) and HEK293T/ACE2/TMPRSS2/CIITA (HEK293T/ATC) cells, respectively (**Table S1B**). Of those, 0.5% (n=119) of A549 and 1.3% (n=389) of HEK293T peptides were derived from SARS-CoV-2 proteins (**Table S1C,D**). This result aligns with the overall low percentage of viral peptides reported in previous studies that investigated the HLA-II immunopeptidome of EBV-B cells infected with measles virus (Ovsyannikova et al., 2003) and Monocyte-derived DCs (moDC) pulsed with recombinant H1-HA protein of influenza virus (Cassotta et al., 2020). Furthermore, the recovered HLA-II peptides exhibited the expected 12-25 amino acid length distribution (**Figure 1D**). We examined the amino acid sequences of detected peptides and checked if these sequences conformed to the binding preferences of the HLA-II alleles expressed in A549 and HEK293T cells. The binding motifs, deconvoluted by Gibbs Cluster (Andreatta et al., 2017), agreed with known preferences of HLA-DR heterodimers (Abelin et al., 2019) expressed in these cell lines (**Figure 1E, Table S1A**). As expected, the deconvoluted motifs matched more to HLA-DR heterodimers, and less so to HLA-DP and HLA-DQ heterodimers, in both cell lines, as HLA-DR is often expressed at higher levels when compared to HLA-DP and HLA-DQ (Taylor et al., 2021). Notably, some of the HLA-II alleles expressed by the two cell lines, and confirmed by motif deconvolution are highly prevalent in the European (EUR) and United States (USA) populations including DRB1*07:01 (∼14% EUR, 12% ∼USA) expressed by A549 and DRB1*15:01 (∼14% EUR, ∼11% USA) and DRB5*01:01 (∼16% EUR) expressed by HEK293T (**Table S1A**).

Finally, we confirmed that CIITA-transduced cells presented peptides derived from extracellular proteins, as should be expected for HLA-II presentation. To this end, we quantified the fraction of HLA-II peptides derived from bovine serum albumin (BSA), a non-human protein present in the cell growth medium, as done previously (Forlani et al., 2021). As a negative control, we examined the HLA-I immunopeptidome of the same cells (Weingarten-Gabbay et al., 2021). Since HLA-I peptides are mostly processed from endogenous proteins, we expected low representation of the exogenous BSA protein in these data. Indeed, we observed 5.2- and 4.6-fold more BSA-derived peptides in the HLA-II peptidome than the HLA-I peptidome in A549/ATC and HEK293T/ATC cells, respectively (**Figure 1G**). Moreover, BSA-derived HLA-II peptides across both A549/ATC and HEK293T/ATC experiments had the expected lengths of 12-25 amino acids (∼87% of peptides), matching the distribution of human-derived HLA-II peptides in the same samples (**Figure 1F**). In contrast, the BSA-derived peptides in HLA-I samples had longer lengths than canonical HLA-I peptides, suggesting that these peptides might have arisen from exogenous peptidase trimming and binding to empty surface HLA-I. Altogether, these data indicate that CIITA-transduced A549/AT and HEK293T/AT cells can present HLA-II peptides generated from endocytosed proteins.

### SARS-CoV-2 HLA-II peptides from canonical and out-of-frame overlapping ORFs

We next analyzed the HLA-II presented peptides derived from SARS-CoV-2 proteins, detecting 469 unique peptides from canonical viral proteins: N, S, M, ORF3a, ORF6, non-structural protein 3 (nsp3) and nsp4 (**Table S1C,D, Figure 2A**). Examining the distribution of HLA-II peptides across SARS-CoV-2 proteins, we observed the expected clustering of peptides into nested sets, with shared core sequences but different N- or C-terminal ends (**Figures 2B** and S2). This pattern is a hallmark of the HLA class II pathway and differentiates it from the class I presentation. While MHC class I molecules structurally constrain the length of loaded peptides, the open structure of the binding groove of MHC-II molecules allows interaction with peptides of variable length, with parts of the peptides protruding out of the binding groove (Lippolis et al., 2002). Interestingly, although A549/ATC and HEK293T/ATC cells express different HLA-II alleles (**Supp Table 1A**) with distinct binding preferences, some clusters contained peptides from both cell lines. This observation suggests that viral antigen processing steps upstream of HLA-II peptide loading play a key role in shaping the HLA-II immunopeptidome.

**Figure 2.**
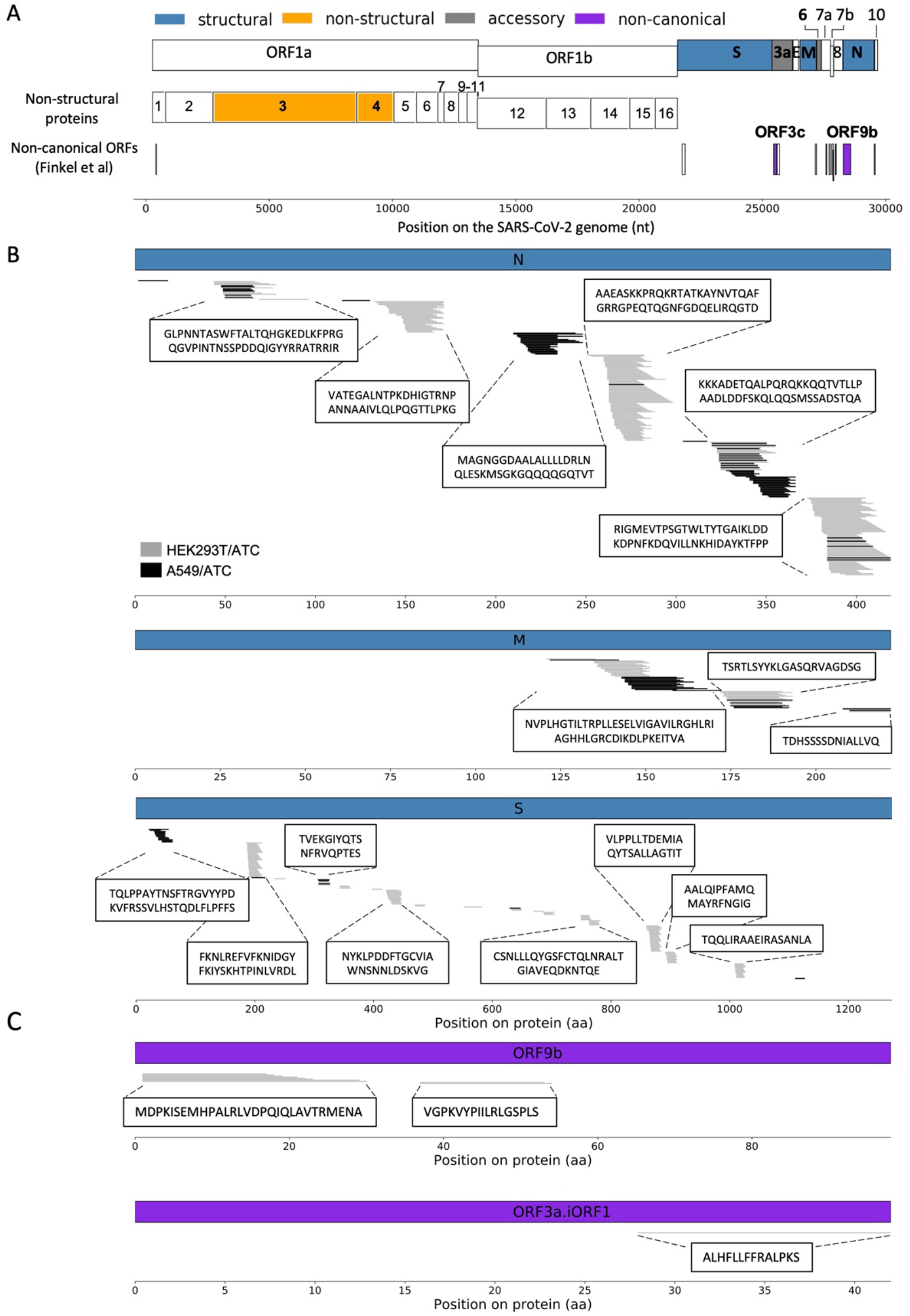
SARS-CoV-2 HLA-II immunopeptidome. **(A)** Summary of viral proteins presented on HLA-II complexes in infected 549/ATC and HEK293T/ATC cells. **(B)** The location of HLA-II peptides in canonical structural proteins M, N and S. Peptides detected in A549/ATC and HEK293T/AT/C cells are depicted in black and gray, respectively. **(C)** The location of HLA-II peptides in two non-canonical internal ORFs: ORF9b and ORF3c.

The untargeted nature of our analysis allowed us to search for peptides originating from non-canonical ORFs (Finkel et al., 2020) in addition to the canonical SARS-CoV-2 proteins. Overall, we detected 11 peptides from two internal overlapping ORFs: ORF9b (overlapping with N, also called N.iORF1) and ORF3c (overlapping with ORF3a, also called 3a.iORF1). ORF9b gave rise to 10 peptides, all of which clustered into two nested sets in the first half of the protein (**Table S1C,D, Figure 2C**). From each nested set, at least one peptide was predicted to bind one of the HLA-II alleles expressed in HEK293T cells: DRB5*01:01 and DQA1*01:02/DQB1*06:02 in the first and second sets, respectively. Interestingly, HLA-II peptides arose from a different region of the ORF9b protein compared to HLA-I peptides, which originated from the C-terminal region (Weingarten-Gabbay et al., 2021). We identified one HLA-II peptide (ALHFLLFFRALPKS) from ORF3c. Although it was only one peptide, it was observed in the same fraction (fxn 3) in two biological replicates and was a high scoring peptide, which supports its authenticity. To the best of our knowledge, this is the first experimental evidence for ORF3c expression at the protein level since it was originally detected using ribosome profiling (Finkel et al., 2020) and computational predictions (Cagliani et al., 2020; Firth, 2020; Jungreis et al., 2021).

### SARS-CoV-2 HLA-II peptides co-localize with epitopes that elicit CD4+ T cell responses in COVID-19 patients

To evaluate if the HLA-II peptides that we detected by mass spectrometry contribute to T cell responses in COVID-19 patients, we compared our HLA-II viral peptides to reported CD4+ epitopes derived from SARS-CoV-2 proteins. We used a curated dataset of T cell epitopes reported by Grifoni et al. (Grifoni et al., 2021). This dataset combines 9 studies that tested CD4+ T cell responses using various assays including ELISpot, Intracellular Cytokine Staining (ICS) and Activation Induced Markers (AIM) (**Figure 3A**) (Keller et al., 2020; Le Bert et al., 2020, 2021; Mahajan et al., 2021; Mateus et al., 2020; Nelde et al., 2021; Peng et al., 2020; Prakash et al., n.d.; Tarke et al., 2021).

**Figure 3.**
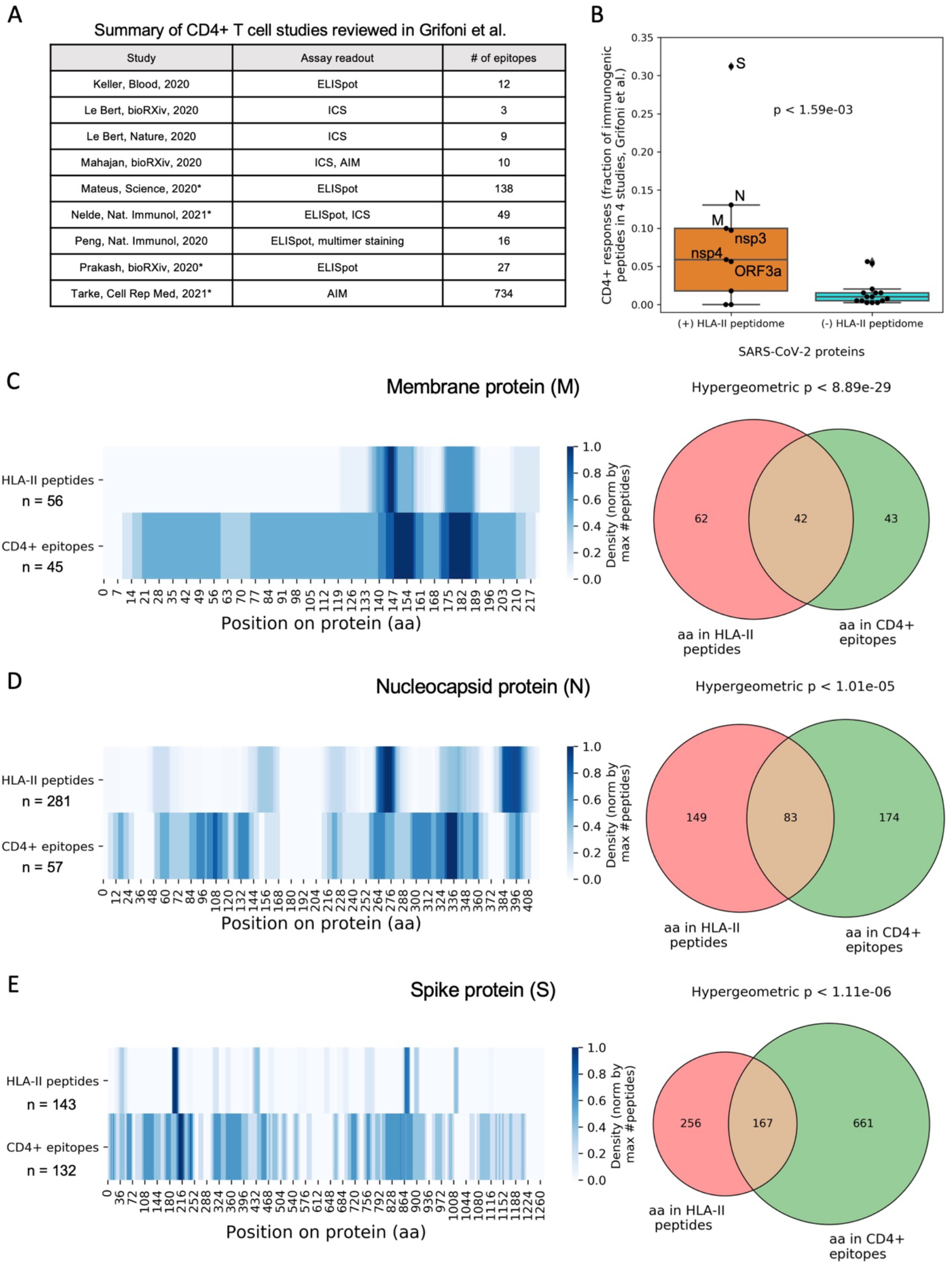
Systematic comparison between the SARS-CoV-2 HLA-II immunopeptidome and known CD4+ T cell epitopes. **(A)** A list of nine studies that identified CD4+ T cells epitopes in COVID-19 patients as described in Table 1 of Grifoni et al (Grifoni et al., 2021). Studies that included peptides from all canonical SARS-CoV-2 proteins are highlighted with an asterisk. **(B)** Comparing the immunogenicity of SARS-CoV-2 proteins that were detected on the HLA-II complex to proteins that were not. For each protein, we plotted the fraction of CD4+ T cell epitopes from all the detected epitopes in four genome-wide studies (denoted with asterisk in panel A). Wilcoxon-rank sum p-value is shown. **(C-E)** Comparing the location of HLA-II peptides that were detected by mass spectrometry in infected A459/AT/CIITA and HEK293T/AT/CIITA cells to previously reported CD4+ T cell epitopes. (left) heatmap showing the density of HLA-II peptides and the reported CD4+ T cell epitopes across individual viral proteins. Rows were normalized according to the maximal density in each row. (right) venn diagram showing the number of amino acids that are covered by HLA-II peptides, CD4+ T cell epitopes and both. To reduce background levels, we counted amino acids with a greater coverage than the mean of each row. Hypergeometric p-value is shown for each viral protein.

First, we checked the overlap between viral proteins that generated HLA-II peptides and those known to contain CD4+ T cell epitopes. We computed the fraction of total CD4+ epitopes that were derived from each of the SARS-CoV-2 proteins. To avoid biases stemming from over-representation of highly characterized viral proteins, such as S, N and M, we limited our analysis to four studies that surveyed the entire canonical SARS-CoV-2 proteome (highlighted in asterisk in **Figure 3A**) (Mateus et al., 2020; Nelde et al., 2021; Prakash et al., n.d.; Tarke et al., 2021). As expected, viral proteins for which we observed HLA-II peptides elicited T cell responses significantly more frequently than those with no detectable HLA-II peptides (Wilcoxon rank-sum p < 10^-3^, **Figure 3B**).

We then considered individual viral proteins and tested if the HLA-II peptides co-localize with regions that elicited CD4+ T cell responses in COVID-19 patients. We focused on the three structural proteins that gave rise to the majority of the detected HLA-II peptides: N (n=281), S (n=143), and M (n=56). These three proteins were also the most abundant source of epitopes in the compiled T cell data, accounting for 54% of the total CD4+ epitopes detected in the canonical SARS-CoV-2 proteome. To determine if HLA-II peptides were derived from regions that were more immunogenic in patients, we counted the number of amino acids that were covered by HLA-II peptides, CD4+ epitopes and both, and computed a hypergeometric p-value that estimates the overlap between these two groups. Since we compared peptide localization within individual proteins, we accounted for epitopes reported in all 9 CD4+ T cell studies listed in **Figure 3A**, including studies that examined only a few SARS-CoV-2 proteins. We found a statistically significant enrichment of HLA-II peptides in regions from which CD4+ epitopes were derived with p<10^-29^, p<10^-5^, and p<10^-5^ for M, N and S, respectively (**Figure 3C-E**).

Furthermore, our analysis showed that HLA-II peptides greatly overlap the two immunodominant regions reported in the M protein. These regions, M:144-163 and M:173-192, were recently identified as hotspots of CD4-restricted epitopes that elicited T cell responses in eight and six COVID-19 convalescent samples, respectively (Keller et al., 2020). The HLA-II peptides that we detected by mass spectrometry were also confined to the same region of M with highest density in the two reported hot spots: M:121-171 (n=34 peptides) and M:172-192 (n=16 peptides), both detected in HEK293T/ATC and A549/ATC cells (**Figure 2B and 3C**, **Table S1C,D,F**). Moreover, the predicted HLA restriction of the two T cell epitopes from M:144-163 described by Keller et al.

(Keller et al., 2020) matches with two of the HLA-II alleles expressed in A549: DRB1*11:04 and DRB4*01:01. Altogether, these results indicate that the HLA-II immunopeptidome of SARS-CoV-2 co-localizes with reported CD4+ T cell epitopes and defines the immunogenic regions in the M protein.

### Distinct subsets of viral proteins are presented on the HLA-I and HLA-II complexes

In the context of vaccine design, it is important to decipher both CD4+ and CD8+ T cell epitopes, since effective vaccines require potent induction of both arms of the adaptive cellular response. Many of the current vaccines strategies rely on the delivery of a single viral protein to invoke both CD8+ and CD4+ T cell responses and mainly focus on the structural proteins S or N (Barouch, 2022; Creech et al., 2021; Dai & Gao, 2021; Krammer, 2020; Kyriakidis et al., 2021; Oronsky et al., 2022; Silva et al., 2022). However, HLA-I and HLA-II presentation, which facilitate CD4+ and CD8+ responses, respectively, have distinct antigen processing steps, raising the likelihood that each of these pathways samples a different subset of viral proteins.

To compare viral proteins presented on HLA-I versus HLA-II complexes, we examined the HLA-I(Weingarten-Gabbay et al., 2021) and HLA-II immunopeptidomes of SARS-CoV-2 infected HEK293T/ATC and A549/ATC cells. We computed the number of peptides observed from each source protein and assessed the representation of proteins in four groups: Non-structural proteins, structural proteins, accessory proteins, and non-canonical ORFs. HLA-II presentation was dominated by the structural proteins N, S, and M, accounting for 95.2% of the detected peptides in HEK293T/ATC and A549/ATC cells, with negligible contribution of the non-structural (1.2%), accessory (1.4%) and non-canonical proteins (2.2%) (**Figure 4A**). In contrast, only 27% of the detected HLA-I peptides were derived from structural proteins, with a large fraction of peptides from non-structural proteins (45.9%) and non-canonical ORFs (24.3%) (**Figure 4B**). Together, these results point to a different presentation profile of SARS-CoV-2 on the HLA-I and HLA-II complexes and suggest that CD4+ and CD8+ T cells engage with different parts of the virus.

**Figure 4.**
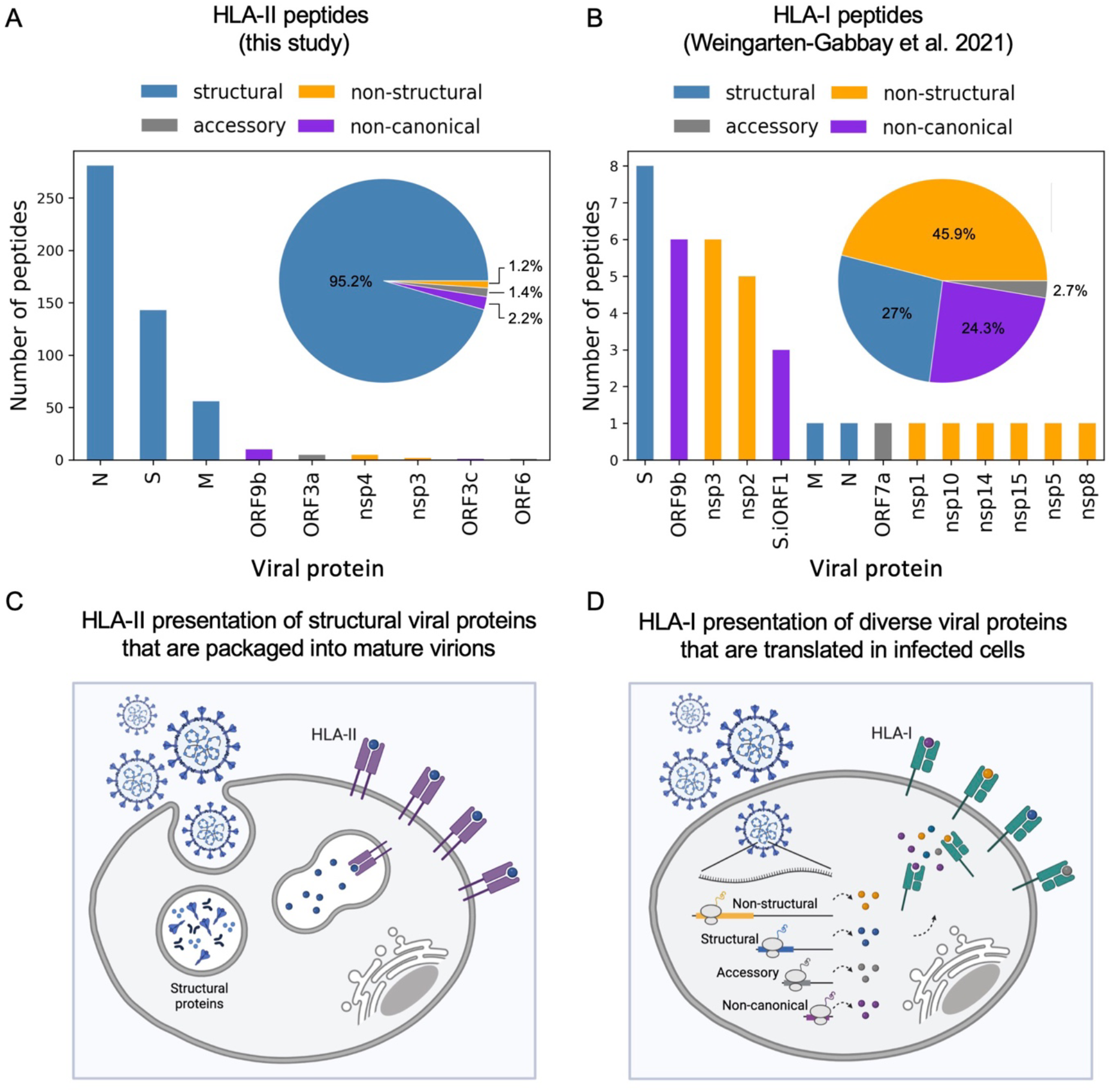
SARS-CoV-2 protein representation on the HLA-I and HLA-II complexes. **(A)** A bar chart showing the number of HLA-II peptides detected in SARS-CoV-2 infected A549/AT/CIITA and HEK293T/AT/CIITA cells for each viral protein. Inside the frame is a pie chart showing the relative abundance of peptides derived from structural proteins, non-structural proteins, accessory proteins and non-canonical ORFs. **(B)** Similar to (A) for HLA-I peptides reported in Weingarten-Gabbay et al. (Weingarten-Gabbay et al., 2021). **(C)** A cartoon illustrating the source proteins for the HLA-II processing and presentation pathway. Viruses are endocytosed by an antigen presenting cell; The viral structural proteins are cleaved within the endosomal–lysosomal compartment and loaded onto HLA-II complexes. **(D)** A cartoon illustrating the source proteins for the HLA-I processing and presentation pathway. Viral proteins are produced from the translation of genomic and subgenomic viral RNAs in infected cells. These proteins are cleaved and loaded onto HLA-I complexes.

## DISCUSSION

We provide the first genomic landscape of SARS-CoV-2 peptides that are naturally processed and presented on the HLA-II complex. This genome-wide view allowed us to systematically compare the HLA-II immunopeptidome to the T cell epitopes in COVID-19 patients and to the HLA-I immunopeptidome of SARS-CoV-2 and to uncover new insights into antigen presentation.

Our work adds to a growing list of studies that employ overexpression of the CIITA master regulator to infer the HLA-II immunopeptidome of cancer cells and viruses (Becerra-Artiles et al., 2019, 2022; Forlani et al., 2021; Hos et al., 2022). Although the cells we profiled were not professional antigen-presenting cells (APCs), we believe that our measurements uncovered genuine HLA-II peptides: The peptides we detected were in the expected length range and contained sequence motifs that matched with the HLA-II alleles of respective cell lines. In addition, we detect presentation of exogenous proteins derived from extracellular bovine serum. Most importantly, our immunopeptidome data successfully captured the SARS-CoV-2 CD4+ T cell epitopes that were detected in a wide range of independent studies using conventional targeted T cell assays. The source proteins of HLA-II peptides in our study were also found to be the most immunogenic proteins in convalescent COVID-19 patients. Moreover, the HLA-II peptides co-localized with regions in the SARS-CoV-2 proteins that are known to elicit strong CD4+ T cell responses.

In the context of highly pathogenic viruses, CIITA induction in HLA-II null cell lines provides an efficient platform to probe the entire viral genome within the restrictions of high-containment facilities. To date, HLA-II immunopeptidome studies of SARS-CoV-2 were limited to a single protein, using pulse experiments with a recombinant protein (Knierman et al., 2020; Parker et al., 2021), or four exogenously expressed proteins using plasmid overexpression (Nagler et al., 2021). By inducing the HLA-II pathway in SARS-CoV-2 infected cells, we could detect HLA-II peptides from the entire viral genome. Although HLA-II peptides are predominantly presented by APCs, such as dendritic cells and macrophages, they can also be presented by non-immune cells upon induction of HLA-II expression, such as lung epithelial cells exposed to IFN-gamma (Neuwelt et al., 2020; Wosen et al., 2018). Thus, the CIITA induction system can also recapitulate naturally occurring HLA-II presentation events of infected cells *in-vivo*. Overexpressing CIITA in the same cell lines in which we profiled the HLA-I immunopeptidome allowed us to use the same reagents and inactivation protocol that we optimized for Biosafety Level 3 (BSL3) laboratory (Weingarten-Gabbay et al., 2022). Thus, we believe that this approach can readily be extended to additional high-containment viruses.

Our analysis uncovers striking differences in the subset of viral proteins that are presented on HLA-I versus HLA-II complexes. We hypothesize that the observed differences stem from the different stages of the viral life cycle at which viral proteins are processed and loaded onto HLA molecules (**Figure 4C,D**). The HLA class II pathway can present peptides from exogenous sources, e.g. mature virions and infected cell debris, that are cleaved within endosomal– lysosomal compartments and loaded onto HLA-II complexes (**Figure 4C**). Thus, the repertoire of HLA-II peptides reflects the viral proteins that are part of mature virus particles. In contrast, the HLA-I pathway samples viral proteins that are actively translated in the cytoplasm of infected cells. Thus, the HLA-I immunopeptidome mirrors the translatome of the virus, including non-structural proteins, accessory proteins, and non-canonical proteins, which are not necessarily incorporated into mature virions (**Figure 4D**). These differences may extend to other viruses as well, many of which encode proteins that are required for viral replication in infected cells but are not packaged into mature virions. The distinct repertoire of viral peptides that are presented on HLA-I versus HLA-II complexes suggests that CD4+ and CD8+ T cells recognize different parts of the viral genome. This observation warrants revision of the current “one-protein-fits-all” approach that forms the basis of most synthetic vaccines, including the broadly administered COVID-19 mRNA vaccines. Rather, incorporating targets from different classes of viral proteins, including structural, non-structural and non-canonical proteins, has the potential to invoke a more holistic response of both CD4+ and CD8+ T cells.

An intriguing question in T cell immunology is what determines the immunodominance of a specific protein or a region within a protein. Observing T cell reactivity in convalescent patients to immunodominant epitopes represents the final outcome in a chain of events that determine which peptides will elicit a T cell response. Some factors defining the immunodominance of a given epitope include protein expression levels, accessibility to proteolytic cleavage and antigen processing, loading onto the HLA complex, the presence of a matching T cell receptor (TCR) in the repertoire of naive T cells, and the binding affinity of an HLA-peptide complex with its matching TCR. Since the immunopeptidome represents antigen processing and presentation steps occurring prior to interaction with TCR, it distinguishes between epitope selectivity at the level of presentation versus T cell recognition. A recent study identified that the HLA-II peptidome of influenza virus defines H1-HA immunodominant regions targeted by memory CD4+ T cells (Cassotta et al., 2020). Similarly to this observation, we found that the two immunodominant hotspots in the M protein (Keller et al., 2020) greatly overlap the HLA-II immunopeptidome of infected cells, suggesting that antigen processing and presentation steps define the immunodominant regions of M. Together, these observations demonstrate the robustness of immunopeptidome analysis for identifying the most relevant protein regions for designing T cell assays and selecting candidates for vaccines.

Our work uncovers HLA-II peptides from two non-canonical overlapping ORFs: ORF9b and ORF3c, and provides the first experimental evidence for ORF3c expression at the peptide level. In contrast to HLA-I presentation, in which non-canonical peptides were enriched on the HLA-I complex, non-canonical peptides represented only a small fraction of the HLA-II immunopeptidome. However, the detection of peptides from non-canonical ORFs in both the HLA-I and the HLA-II immunopeptidomes emphasizes the importance of incorporating non-canonical ORFs into T cell studies to achieve a complete picture of the antiviral immune profile. The discovery of a peptide from ORF3c highlights another important aspect of immunopeptidome studies and its potential impact on our understanding of non-canonical ORFs. Although non-canonical ORFs have numerous roles in the viral life cycle, including regulating viral gene expression and modulating virus infection, their detection in tryptic proteome experiments is often challenging due to their small size, shorter half-life, and in some cases, the lack of observable tryptic peptides. Of the 23 non-canonical ORFs that were identified in the SARS-CoV-2 genome (Finkel et al., 2020), only one (ORF9b) has so far been detected in global tryptic proteomic experiments (Weingarten-Gabbay et al., 2021). The longer half-life of the HLA peptide complex compared to the non-canonical ORF translation product in the cell may increase the probability of detection by mass spectrometry (Ruiz Cuevas et al., 2021). As an example, although we and others identified three peptides from S.iORF1 (an internal overlapping ORF in spike) on the HLA-I complex (Nagler et al., 2021; Weingarten-Gabbay et al., 2021), this protein was not detected in whole tryptic proteome analysis of the same infected cell lysates. ORF3c was recently shown to inhibit innate immunity by restricting IFN-β production, exposing an important mechanism of SARS-CoV-2 immune evasion (Stewart et al., 2022). Our study provides evidence that this important non-canonical protein is expressed in cells that are infected with SARS-CoV-2. Thus, in addition to enhancing our understanding of viral antigen presentation, immunopeptidome studies contribute to our basic understanding of viruses by illuminating the complete set of canonical and non-canonical viral proteins.

There are limitations to this study. Although CIITA overexpression activated the HLA-II pathway in the cell lines used in our study, allowing the investigation of SARS-CoV-2 HLA-II immunopeptidome, these cells may not capture the unique biology of how APCs subtypes uptake and process mature virions and virus infected cells *in vivo*. In contrast to the cells profiled in this study, APCs are not naturally infected by SARS-CoV-2. It is possible that productive infection impacts the antigen processing and presentation steps and provides an internal source of viral proteins, in addition to the endocytosed particles, for production of HLA-II peptides. For instance, Ghosh and colleagues (Ghosh et al. 2020) showed that β-Coronaviruses can traffic to lysosomes and egress by Arl8b-dependent lysosomal exocytosis. In this context, lysosomes are deacidified, which can inactivate proteolytic enzymes in infected cells and impair antigen presentation. Further, our study only uncovered the peptides that are presented by the HLA-II alleles expressed in A549 and HEK293T cells. As such, we might have missed SARS-CoV-2 HLA-II peptides that were incompatible with A549 and HEK293T cells. Nonetheless, our study identifies HLA-II peptides that can be presented on virally infected cells and provides valuable insights into which viral proteins are more likely to be presented by SARS-CoV-2 infected cells.

## METHODS

### Plasmid

Lentiviral vectors pLOC_hACE2_PuroR and pLOC_hTMPRSS2_BlastR, harboring human ACE2 and TMPRSS2, respectively, have been described (Chen et al., JVI). To generate a lentiviral vector containing human CIITA, we amplified the CIITA cDNA from pcDNA3 myc CIITA (Addgene #14650) and cloned it into pTRIP-SFFV-Hygro-2A (previously described(Gentili et al., 2023)) via Gibson assembly. The resultant plasmid was named pTRIP-SFFV-Hygro-2A-myc-CIITA.

### Cell culture

Human embryonic kidney HEK293T cells (female), human lung A549 cells (male), and African green monkey kidney epithelial Vero E6 cells (female) were maintained at 37°C and 5% CO2 in DMEM containing 10% FBS. To generate HEK293T and A549 cells overexpressing human ACE2, TMPRSS2, and CIITA, we transduced these cells with lentiviral vectors pLOC_hACE2_PuroR, pLOC_hTMPRSS2_BlastR, and pTRIP-SFFV-Hygro-2A-myc-CIITA, and selected for the triple-transduced cells in culture medium supplemented with 1 µg/ml each of puromycin and blasticidin and 320 µg/ml of hygromycin. A375 cells were obtained from ATCC (ATCC ® CRL-1619). A375 cells were grown in ATCC-formulated Dulbecco’s Modified Eagle’s Medium (Catalog No. 30-2002) with fetal bovine serum to a final concentration of 10% using ATCC guidelines. A375 cells were harvested by trypsinization (Trypsin-EDTA 0.25%, Gibco™ 25200056), pelleted and rinsed in PBS twice. Pellets were snap frozen and stored at −80 °C.

### Flow cytometry

1.5*10^6 A549 or A375 cells were incubated with 2ul FITC anti HLA-DR, DP, DQ antibody (BD Pharmingen #562008) in 100ul PBS at 4°C for 45 minutes. Cells were washed three times with PBS, resuspended in 400ul PBS and analyzed using a CytoFLEX flow cytometer.

### SARS-CoV-2 virus stock preparation and titration

The SARS-CoV-2 USA-WA1/2020 isolate (NCBI accession number: MT246667) was deposited by the Centers for Disease Control and Prevention and obtained through BEI Resources, NIAID, NIH (NR-52281). We then passaged the virus twice onto Vero E6 cells to obtain the P2 stock, as previously described (Chen et al., JVI). The virus titration was performed on Vero E6 cells. All experiments in this study utilized the P2 stock.

### Quantification of virus infectivity using immunofluorescence

A549 and 293T cells stably overexpressing ACE2, TMPRSS2, and CIITA were infected with SARS-CoV-2 at an MOI of 0.5, 1, or 3 for 12, 18, 24, 36, or 48 hours. At indicated times, the culture medium was removed, and the cells were fixed with 4% paraformaldehyde for 60 minutes at room temperature. The cells were then permeabilized with 0.1% of Triton X-100 in PBS for 10 minutes and hybridized with anti-SARS-CoV nucleocapsid (rabbit polyclonal) antibody (1:2000, Rockland, #200-401-A50) at 4°C overnight. AlexaFluor 568 goat anti-rabbit antibody (Invitrogen, #A11011) was used as the secondary antibody. Finally, DAPI was used to stain cell nuclei. Images were captured with an EVOS microscope using a 10x lens, and the percentage of infected cells was calculated with ImageJ.

### Immunoprecipitation of HLA-II complexes from cells

Cells from 3×15cm dishes were scraped into 2.5ml/dish of cold lysis buffer (20mM Tris, pH 8.0, 100mM NaCl, 6mM MgCl2, 1mM EDTA, 60mM Octyl β-d-glucopyranoside, 0.2mM Iodoacetamide, 1.5% Triton X-100, 50xC0mplete Protease Inhibitor Tablet-EDTA free and PMSF) obtaining a total of ∼9 mL lysate. This lysate was split into 6 eppendorf tubes, with each tube receiving 1.5 mL volume, and incubated on ice for 15 min with 1ul of Benzonase (Thomas Scientific, E1014-25KU) to degrade nucleic acid. The lysates were then centrifuged at 4,000 rpm for 22 min at 4°C and the supernatants were transferred to another set of 6 eppendorf tubes containing a mixture of pre-washed beads (Millipore Sigma, GE17-0886-01) and 12.5 uL (12.5 ug) of MHC class II antibodies in a 3:1:1 mixture of TAL-1B5 (Abcam, ab20181), EPR11226 (Abcam, ab157210) and B-K27 (Abcam, ab47342). The immune complexes were captured on the beads by incubating on a rotor at 4°C for 3hr in the BSL3 lab. Virus inactivation was confirmed before subsequent samples processing outside the BSL3 using plaque assay (Weingarten-Gabbay et al., 2021, 2022). In total, nine washing steps were performed; one wash with 1mL of cold lysis wash buffer (20mM Tris, pH 8.0, 100mM NaCl, 6mM MgCl2, 1mM EDTA, 60mM Octyl β-d-glucopyranoside, 0.2mM Iodoacetamide, 1.5% Triton X-100), four washes with 1mL of cold complete wash buffer (20mM Tris, pH 8.0, 100mM NaCl, 1mM EDTA, 60mM Octyl β-d-glucopyranoside, 0.2mM Iodoacetamide), and four washes with 20mM Tris pH 8.0 buffer. Dry beads were stored at −80°C until mass-spectrometry analysis was performed.

### HLA-II peptidome desalting and LC-MS/MS data generation

HLA peptides were eluted and desalted from beads as follows: wells of the tC18 40mg Sep-Pak desalting plate (Waters, Milford, MA) were activated with 2x 1 mL of methanol (MeOH) and 500 µL of 99.9% acetonitrile (ACN)/0.1% formic acid (FA), then washed with 4x 1 mL of 1% FA. A 10μmPE fritted filter plate containing the HLA-IP beads was placed on top of the Sep-Pak plate. To dissociate peptides from HLA molecules and facilitate peptides binding to the tC18 solid phase, 200 µL of 3% ACN/5% FA was added to the beads in the filter plate. 100 fmol internal retention time (iRT) standards (Biognosys SKU: Ki-3002-2) was spiked into each sample as a loading control and pushed through both the filter plate and 40 mg Sep-Pak plate. Following sample loading there was one wash with 400 µL of 1% FA. Beads were then incubated with 500 µL of 10% acetic acid (AcOH) three times for 5 min to further dissociate bound peptides from the HLA molecules. The beads were rinsed once with 1 mL 1% FA and the filter plate was removed. The Sep-Pak desalt plate was rinsed with 1 mL 1% FA an additional three times. The peptides were eluted from the Sep-Pak desalt plate using 250 µL of 15% ACN/1% FA and 2x 250 µL of 50% ACN/1% FA. HLA peptides were eluted into 1.5 mL micro tubes (Sarstedt, Nümbrecht, Germany), frozen, and dried down via vacuum centrifugation. Dried peptides were stored at −80°C until microscaled basic reverse phase separation.

Briefly, peptides were loaded on Stage-tips with 2 punches of SDB-XC material (Empore 3M). HLA-II peptides were eluted in three fractions with increasing concentrations of ACN (5%, 15%, and 40% in 0.1% NH4OH, pH 10). Peptides were reconstituted in 3% ACN/5% FA prior to loading onto an analytical column (35 cm, 1.9 µm C18 (Dr. Maisch HPLC GmbH), packed in-house PicoFrit 75 µm inner diameter, 10 µm emitter (New Objective)). Peptides were eluted with a linear gradient (EasyNanoLC 1200, Thermo Fisher Scientific) ranging from 6–30% Solvent B (0.1% FA in 90% ACN) over 84 min, 30–90% B over 9 min and held at 90% B for 5 min at 200 nl/min. MS/MS data were acquired on a Thermo Orbitrap Exploris 480 equipped with (HLA-I) and without (HLA-II) FAIMS (Thermo Fisher Scientific) in data-dependent acquisition. FAIMS compensation voltages (CVs) were set to −50 and −70 with a cycle time of 1.5 s per FAIMS experiment. MS2 fill time was set to 100 ms; collision energy was 30, 34 or 36 CE see **Table S2** for file mappings and LC-MS/MS acquisition variations.

### Proteome analysis from HLA enrichment flow-through

Flow-throughs of the HLA-II IP that were stored as flash-frozen native protein lysates were briefly thawed on ice for ∼15 min. Once thawed, 10% SDS was added for a final concentration of 5% SDS to denature the lysate, resulting in a final volume of ∼1.5 mL lysate which was prepared for S-Trap digestion (Abelin et al., 2023).

Protein concentration was estimated using a BCA assay for scaling of digestion enzymes. Disulfide bonds were reduced in 5 mM DTT for 30 min at 25°C and 1000 rpm shaking and cysteine residues were alkylated in 10 mM IAA in the dark for 45 min at 25°C and 1000 rpm shaking. Lysates were then transferred to a 15 mL conical tube to prepare for protein precipitation. 27% phosphoric acid was added at a 1:10 ratio of lysate volume to acidify and proteins were precipitated with 6x sample volume of ice cold S-Trap buffer (90% methanol, 100 mM TEAB). The precipitate was transferred in successive loads of 3 mL to a S-Trap Midi (Protifi) and loaded with 1 min centrifugation at 4000 x g, mixing the remaining precipitate thoroughly between transfers. The precipitated proteins were washed 4x with 3 mL S-Trap buffer at 4000 x g for 1 min. To digest the deposited protein material, 350 µL digestion buffer (50 mM TEAB) containing both trypsin and endopeptidase C (LysC), each at 1:50 enzyme:substrate, was passed through each S-Trap column with 1 min centrifugation at 4000 x g. The digestion buffer was then added back atop the S-Trap and the cartridges were left capped overnight at 25°C.

Peptide digests were eluted from the S-Trap, first with 500 µL 50 mM TEAB and next with 500 µL 0.1% FA, each for 30 sec at 1000 x g. The final elution of 500 µL 50% ACN/0.1% FA was centrifuged for 1 min at 4000 x g to clear the cartridge. Peptide concentration of the pooled elutions was estimated with a BCA assay, and 10 µg peptide was used for stagetip fractionation. Each 25ug proteome sample was reconstituted in 4.5 mM ammonium formate (pH 10) in 2% (vol/vol) acetonitrile and separated into four fractions using basic reversed phase fractionation on a C-18 Stage-tip. Fractions were eluted at 5%, 12.5%, 15%, and 50% ACN/4.5 mM ammonium formate buffer (pH 10) and dried. Fractions were reconstituted in 3%ACN/5%FA, and 1 ug was used for LC-MS/MS analysis.

Data-dependent acquisition was performed using Orbitrap Exploris 480 V2.0 software in positive ion mode at a spray voltage of 1.8 kV. MS1 spectra were measured with a resolution of 60,000, a normalized AGC target of 300% for, a maximum injection time of 10 ms, and a mass range from 350 to 1800 m/z. The data-dependent mode cycle was set to trigger MS/MS on up to the top 20 most abundant precursors per cycle at an MS2 resolution of 45,000, an AGC target of 30%, an isolation window of 0.7 m/z, a maximum injection time of 105 ms for proteome, and an HCD collision energy of 34%. Peptides that triggered MS/MS scans were dynamically excluded from further MS/MS scans for 20 s in proteome/phosphoproteome/ubiquitylome and for 30 s in acetylome, with a ±10 ppm mass tolerance. Theoretical precursor envelope fit filter was enabled with a fit threshold of 50% and window of 1.2 m/z. Monoisotopic peak determination was set to peptide and charge state screening was enabled to only include precursor charge states 2–6 with an intensity threshold of 5.0e3. Advanced peak determination (APD) was enabled. “Perform dependent scan on single charge state per precursor only” was disabled.

### LC-MS/MS data interpretation

Peptide sequences were interpreted from MS/MS spectra using Spectrum Mill (SM) v 7.08 (proteomics.broadinstitute.org) against a RefSeq-based sequence database containing 41,457 proteins mapped to the human reference genome (hg38) obtained via the UCSC Table Browser (https://genome.ucsc.edu/cgi-bin/hgTables) on June 29, 2018, with the addition of 13 proteins encoded in the human mitochondrial genome, 264 common laboratory contaminant proteins, 553 human non-canonical small open reading frames, 28 SARS-CoV2 proteins obtained from RefSeq derived from the original Wuhan-Hu-1 China isolate NC_045512.2 (https://www.ncbi.nlm.nih.gov/nuccore/1798174254;(Wu et al., 2020)), and 23 novel unannotated virus ORFs whose translation is supported by Ribo-seq(Finkel et al., 2020) for a total of 42,337 proteins. Among the 28 annotated SARS-CoV2 proteins we opted to omit the full-length polyproteins ORF1a and ORF1ab, to simplify peptide-to-protein assignment, and instead represented ORF1ab as the mature 16 individual non-structural proteins that result from proteolytic processing of the 1a and 1ab polyproteins. We added the D614G variant of the SARS-Cov2 Spike protein that is commonly observed in European and American virus isolates, and also added 2036 entries from 6-frame translation of the SARS-Cov2 genome for all possible ORFs longer than 6 amino acids.

Parameters for the SM MS/MS search module for HLA-II immunopeptidomes included: no enzyme specificity; precursor and product mass tolerance of ±10 ppm; minimum matched peak intensity of 30%; ESI-QEXACTIVE-HCD-HLA-v3 scoring; fixed modification: carbamidomethylation of cysteine; variable modifications: cysteinylation of cysteine, oxidation of methionine, deamidation of asparagine, acetylation of protein N-termini, and pyroglutamic acid at peptide N-terminal glutamine; and precursor mass shift range of −18 to 81 Da. For tryptic proteomes, parameters included: ‘‘trypsin allow P’’ enzyme specificity with up to 4 missed cleavages, precursor and product mass tolerance of ± 20 ppm, and 30% minimum matched peak intensity (40% for acetylome). Scoring parameters were ESI-QEXACTIVE-HCD-v2. Allowed fixed modifications included carbamidomethylation of cysteine and selenocysteine.Allowed variable modifications for whole proteome datasets were acetylation of protein N-termini, oxidized methionine, deamidation of asparagine, hydroxylation of proline in PG motifs, pyro-glutamic acid at peptide N-terminal glutamine, and pyro-carbamidomethylation at peptide N-terminal cysteine with a precursor MH+ shift range of −18 to 97 Da.

## ACKNOWLEDGEMENT

We thank Susan Klaeger and members of the Rice lab for valuable discussions. The following reagent was deposited by the Centers for Disease Control and Prevention and obtained through BEI Resources, NIAID, NIH: SARS-Related Coronavirus 2, Isolate hCoV-19/USA-WA1/2020, NR-52281. This study was supported in part by grants from the National Institute of Allergy and Infectious Diseases (U19AI110818 to P.C.S.) and the United States Department of Agriculture (58-3022-2-031 to P.C.S). This work was supported by the National Cancer Institute (NCI) grants U24CA271075, U24CA270823, U24CA210986, and U24CA210979 to S.A.C. and Dr. Miriam and Sheldon G. Adelson Medical Research Foundation to S.A.C. S.W.-G. is the recipient of a Human Frontier Science Program fellowship (LT-000396/2018), EMBO non-stipendiary long-term fellowship (ALTF 883-2017), the Gruss-Lipper postdoctoral fellowship, the Zuckerman STEM Leadership Program fellowship, and the Rothschild Postdoctoral Fellowship. M.S. is supported by Boston University startup funds.

## AUTHORS CONTRIBUTION

S.W.-G. and J.G.A. conceptualized the study. S.W.-G., J.G.A., and M.S. designed the experiments. D.-Y.C., H.B.T., M.G., L.R.P., M.R.B. performed experiments. S.W.-G., S.S., H.B.T., K.R.C., and J.G.A. performed data analysis. N.H., S.A.C., M.S., and P.C.S. supervised the work. S.W.-G., J.G.A., M.S., and P.C.S. wrote the manuscript with contributions from all the authors.

## SUPPPLEMENTAL FIGURES

**Figure S1.**
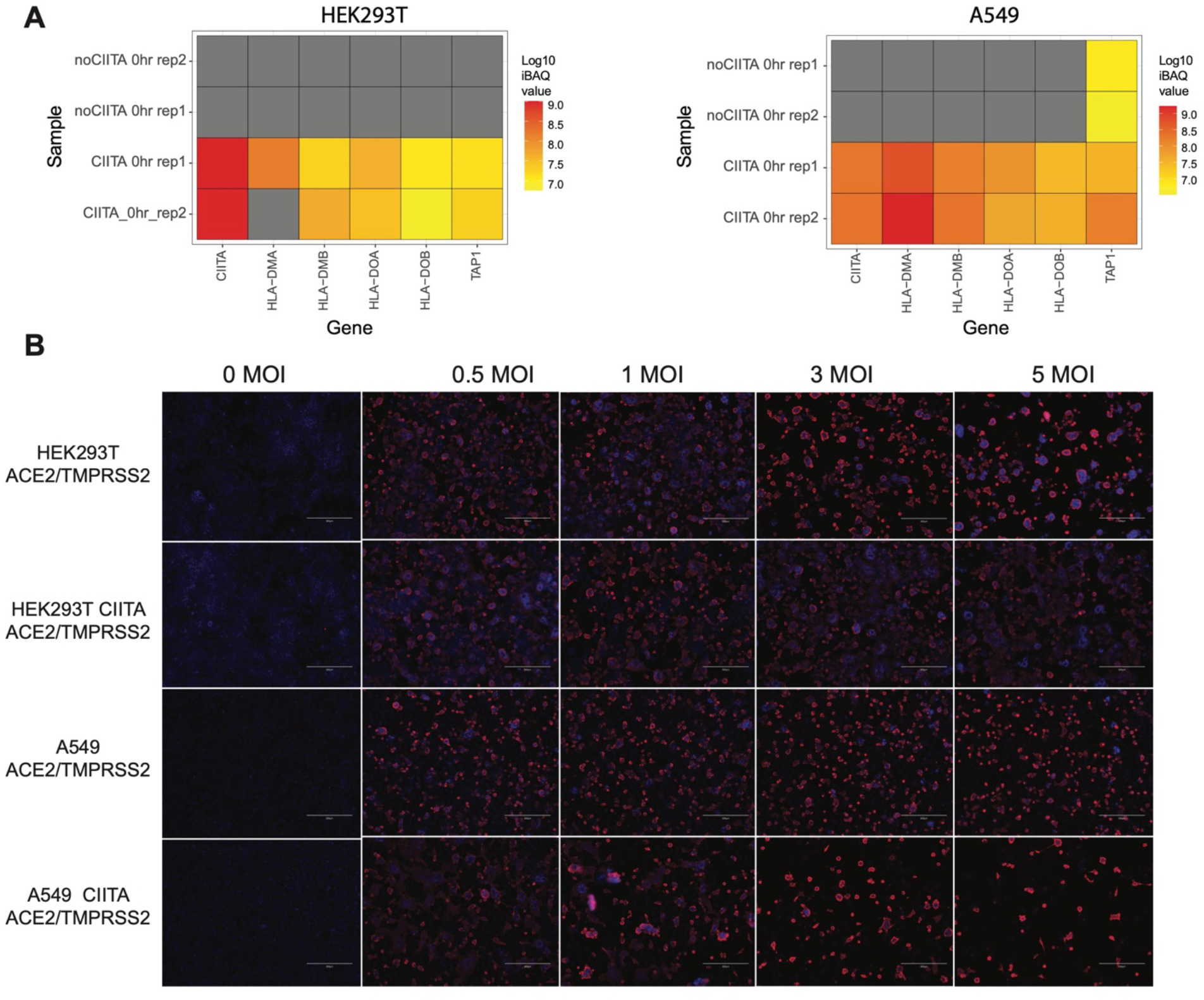
Proper induction of the MHC-II locus and SARS-CoV-2 infectivity in CIITA overexpressing cells. **(A)** Heatmap of log10 iBAQ values for key proteins in the MHC-II locus, observed in whole proteome datasets of A549/AT and HEK293T/AT cells with and without CIITA transduction. Shown are two biological replicates for each condition. **(B)** SARS-CoV-2 infectivity assay comparing A549/AT and HEK293T/AT with and without CIITA transduction. Representative immunofluorescence images at 24 hpi. Red, nucleocapsid; Blue, DAPI. Images were captured with an EVOS microscope using a 10x objective lens.

**Figure S2.**
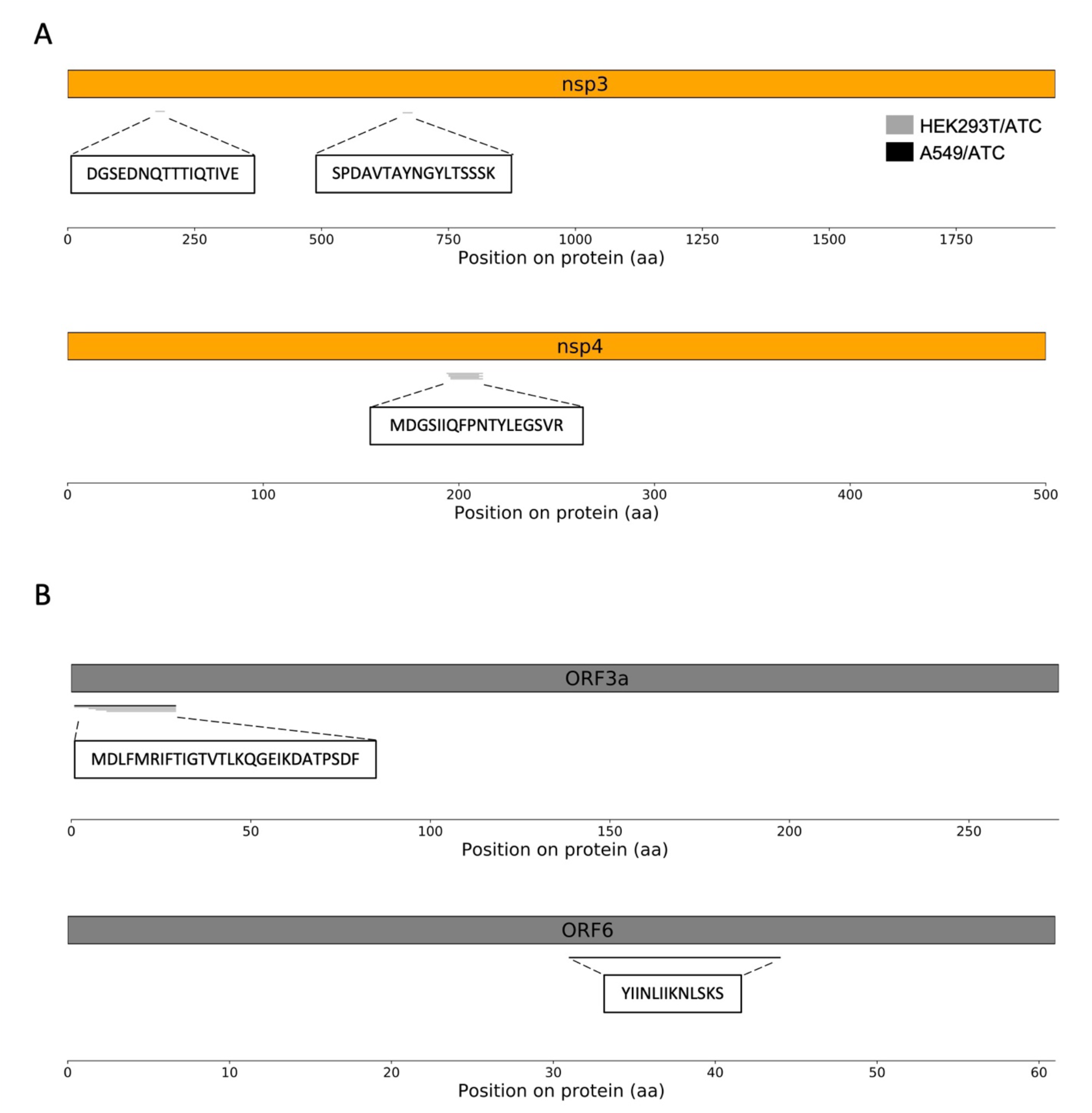
HLA-II peptides in SARS-CoV-2 non-structural and accessory proteins. **(A)** The location of HLA-II peptides across the non-structural proteins nsp3 and nsp4. **(B)** The location of HLA-II peptides across the accessory proteins ORF3a and ORF6. Peptides detected in A549/ATC and HEK293T/ATC cells are depicted in black and gray, respectively.

